# Autophagy promotes cell and organismal survival by maintaining NAD(H) pools

**DOI:** 10.1101/2020.01.31.928424

**Authors:** Lucia Sedlackova, Elsje G. Otten, Filippo Scialo, David Shapira, Tetsushi Kataura, Bernadette Carroll, Elena Seranova, Yoana Rabanal-Ruiz, George Kelly, Rhoda Stefanatos, Glyn Nelson, Francesca Urselli, Animesh Acharjee, Niall Kenneth, Sergey Trushin, Tong Zhang, Charles C. Bascom, Ryan Tasseff, Robert J. Isfort, John E. Oblong, Eugenia Trushina, Masaya Imoto, Shinji Saiki, Michael Lazarou, Manolis Papamichos Chronakis, Oliver D.K. Maddocks, Sovan Sarkar, Alberto Sanz, Viktor I. Korolchuk

## Abstract

Autophagy is an essential catabolic process that promotes clearance of surplus or damaged intracellular components^1^. As a recycling process, autophagy is also important for the maintenance of cellular metabolites during periods of starvation^2^. Loss of autophagy is sufficient to cause cell death in animal models and is likely to contribute to tissue degeneration in a number of human diseases including neurodegenerative and lysosomal storage disorders^3–7^. However, it remains unclear which of the many cellular functions of autophagy primarily underlies its role in cell survival. Here we have identified a critical role of autophagy in the maintenance of nicotinamide adenine dinucleotide (NAD^+^/NADH) levels. In respiring cells, loss of autophagy caused NAD(H) depletion resulting in mitochondrial membrane depolarisation and cell death. We also found that maintenance of NAD(H) is an evolutionary conserved function of autophagy from yeast to human cells. Importantly, cell death and reduced viability of autophagy-deficient animal models can be partially reversed by supplementation with an NAD(H) precursor. Our study provides a mechanistic link between autophagy and NAD(H) metabolism and suggests that boosting NAD(H) levels may be an effective intervention strategy to prevent cell death and tissue degeneration in human diseases associated with autophagy dysfunction.

---

Macroautophagy, hereinafter autophagy, is a cellular trafficking pathway mediated by the formation of double-membraned vesicles called autophagosomes, which ultimately fuse with lysosomes, where their cargo is degraded. By sequestering and clearing dysfunctional cellular components, such as protein aggregates and damaged organelles, autophagy maintains cellular homeostasis whilst also providing metabolites and energy during periods of starvation. Studies using a range of laboratory models from yeast to mammals have established that autophagy is essential for cellular and organismal survival. For example, inducible knockout of core autophagy genes, such as *Atg5,* results in cell death and tissue degeneration in adult mice^3, 8, 9^. However, autophagy-deficient cells such as *Atg5*^−/−^ mouse embryonic fibroblasts (MEFs) are viable in cell culture, which hinders *in vitro* studies of the mechanisms leading to cell death^8–10^. We hypothesized that this apparent discrepancy between the requirement for functional autophagy *in vivo* and *in vitro* could be due to a metabolic shift from oxidative phosphorylation (OXPHOS) to glycolysis. Indeed, whilst differentiated cells with high energy demand, such as neurons, rely on aerobic ATP generation via OXPHOS, the abundance of glucose in standard cell culture conditions allows cells to generate sufficient levels of ATP via glycolysis. This decreased reliance on mitochondrial respiration could then mask an underlying viability defect^11^.

A well-established strategy to reverse cellular reliance on energy generation via aerobic glycolysis and promote mitochondrial OXPHOS, is to replace glucose, the major carbon source in tissue culture media, with galactose^12–16^. Therefore, to investigate whether driving mitochondrial respiration *in vitro* could expose a viability defect in autophagy-deficient cells, we cultured wild-type and *Atg5*^−/−^ MEFs in galactose media. Strikingly, *Atg5*^−/−^ MEFs displayed a rapidly apoptotic phenotype, which was rescued by re-expression of Atg5 (Fig. 1a-c and Extended Data Fig. 1a). Similarly, CRISPR/Cas9 deletion of essential autophagy genes *Atg5*, *Atg7* and *Rb1cc1*, (homologue of human *FIP200*), as well as the loss of lysosomal cholesterol transporter Npc1 required for efficient autophagy^6^, also led to apoptosis upon cell culture in galactose media (Extended Data Fig. 1b, c).

**Fig. 1:**
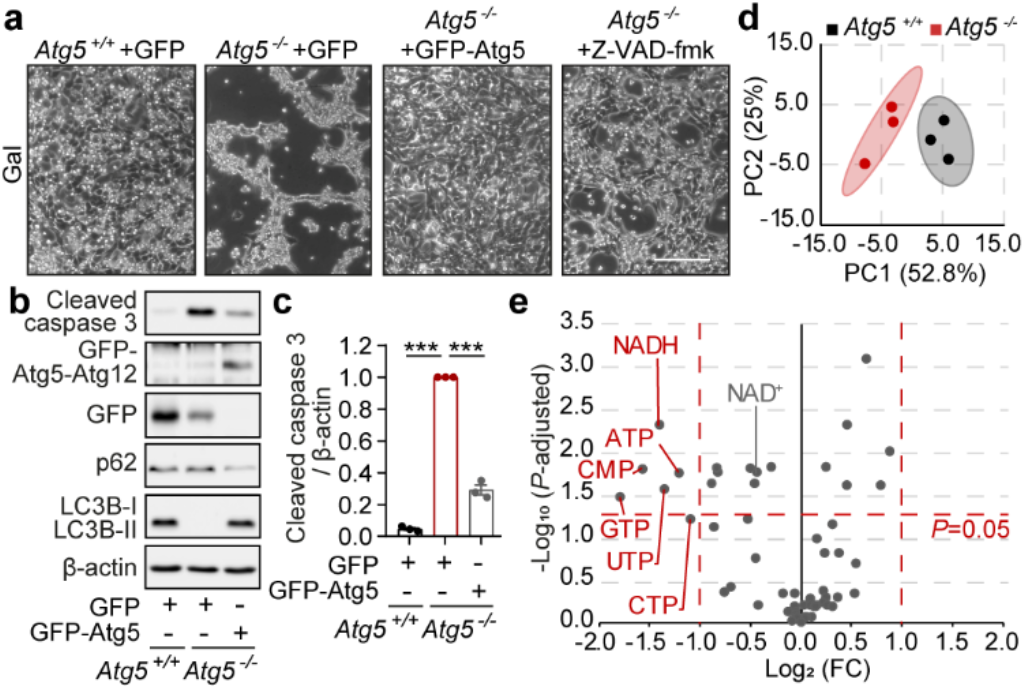
Culture of autophagy-deficient cells in galactose medium results in apoptotic cell death and depletion of NAD(H). **a**, Phase contrast images of *Atg5*^+/+^+GFP, *Atg5*^−/−^+GFP, *Atg5*^−/−^+GFP-Atg5 MEFs, or *Atg5*^−/−^ MEFs supplemented with 20 μM Z-VAD-fmk, cultured in a galactose (gal) medium for 24 h. **b**, **c**, Immunoblot analyses of *Atg5*^+/+^+GFP, *Atg5*^−/−^+GFP, *Atg5*^−/−^+GFP-Atg5 MEFs using microtubule-associated protein 1A/1B light chain 3B (MAP1LC3B, LC3B), p62, cleaved caspase 3 and β-actin antibodies in the same conditions as (**a**). **d**, Two-dimensional principal component analysis (PCA) scores plot of metabolites in *Atg5*^+/+^ (black) vs *Atg5*^−/−^ (red) MEFs cultured in gal medium. **e**, Volcano plot representation of all analysed metabolites in a pairwise comparison of *Atg5*^−/−^ to *Atg5*^+/+^ MEFs. The data were plotted as Log_2_ fold change (FC) versus –Log_10_ of the false discovery rate (FDR) adjusted *P* value. The significance cut-off was set to an adjusted *P* value of 0.05 (−Log_10_(*P*-adjusted)>1,3) and a two-fold change (−1≥Log_2_(FC)≥1). Thresholds are shown as dashed red lines. Data (**c**) are mean ± s.e.m. *P* values were calculated by unpaired two-tailed Student’s *t*-test (**c**) and the multiple *t*-test original FDR method of Benjamini and Hochberg (**e**) on three independent experiments. \*\*\**P*<0.001. Scale bar, 200 μm (**a**).

The rapid nature of cell death suggested an underlying metabolic collapse in autophagy-deficient cells^17^. Loss of autophagy was previously linked to the depletion of numerous cellular metabolites including amino acids, fatty acids and nucleotides, although the relevance of this metabolic defect to the *in vivo* cell death phenotype remains unknown^2^. To investigate the potential metabolic basis of cell death due to autophagy deficiency, we performed an unbiased metabolomics profiling of wild-type and *Atg5*^−/−^ MEFs prior to the onset of cell death. In agreement with a previously proposed general defect in nucleic acid recycling in autophagy-deficient cells^18^, our profiling detected a number of nucleotides that were significantly depleted in *Atg5*^−/−^ MEFs (Fig. 1d, e and Extended Data Fig. 1d). By plotting the magnitude of change against the measure of significance, we identified that the reduced form of nicotinamide adenine dinucleotide (NADH) was the most significantly affected nucleotide in autophagy-deficient cells (Fig. 1e). Importantly, NAD^+^, an oxidised form of the NAD nucleotide, was also significantly decreased, thus suggesting that autophagy-deficient cells present with a depletion of the total pool of the NAD(H) dinucleotide, rather than just a shift in NAD redox state (Fig. 1e).

To test if depletion of NAD(H) is sufficient to decrease cell viability in OXPHOS-dependent cells, we treated wild-type MEFs with FK866, an inhibitor of nicotinamide phosphoribosyltransferase (NAMPT) involved in NAD biosynthesis^19^. Consistent with the requirement of NAD(H) for the survival of OXPHOS-dependent cells, FK866 triggered apoptosis of wild-type MEFs in galactose, but not glucose, media (Fig. 2a-c). We then investigated whether depletion of the total NAD(H) pool in *Atg5*^−/−^ MEFs may be caused by enzymes that use NAD^+^ as a cofactor and cleave it into ADP-ribose and nicotinamide (NAM)^20^. Two main classes of such enzymes, sirtuins (SIRTs) and poly-ADP-ribose polymerases (PARPs), are activated in response to oxidative stress or DNA damage, which could occur in conditions of autophagy deficiency^21, 22^. While short-term activation of PARPs and SIRTs in response to stress is beneficial for adaptation to metabolic or genetic aberrations, uncontrolled NAD^+^ cleavage can contribute to total NAD(H) exhaustion and loss of cell viability^23^. Indeed, we detected increased levels of both, SIRT-dependent histone deacetylase activity and PARP-mediated poly-ADP-ribosylation (PARylation), in *Atg5*^−/−^ MEFs (Extended Data Fig. 2a, b). Crucially, pharmacological inhibition of either class of enzymes partially rescued both, intracellular NAD(H) levels and cell viability of *Atg5*^−/−^ MEFs (Fig. 2d-f).

**Fig. 2:**
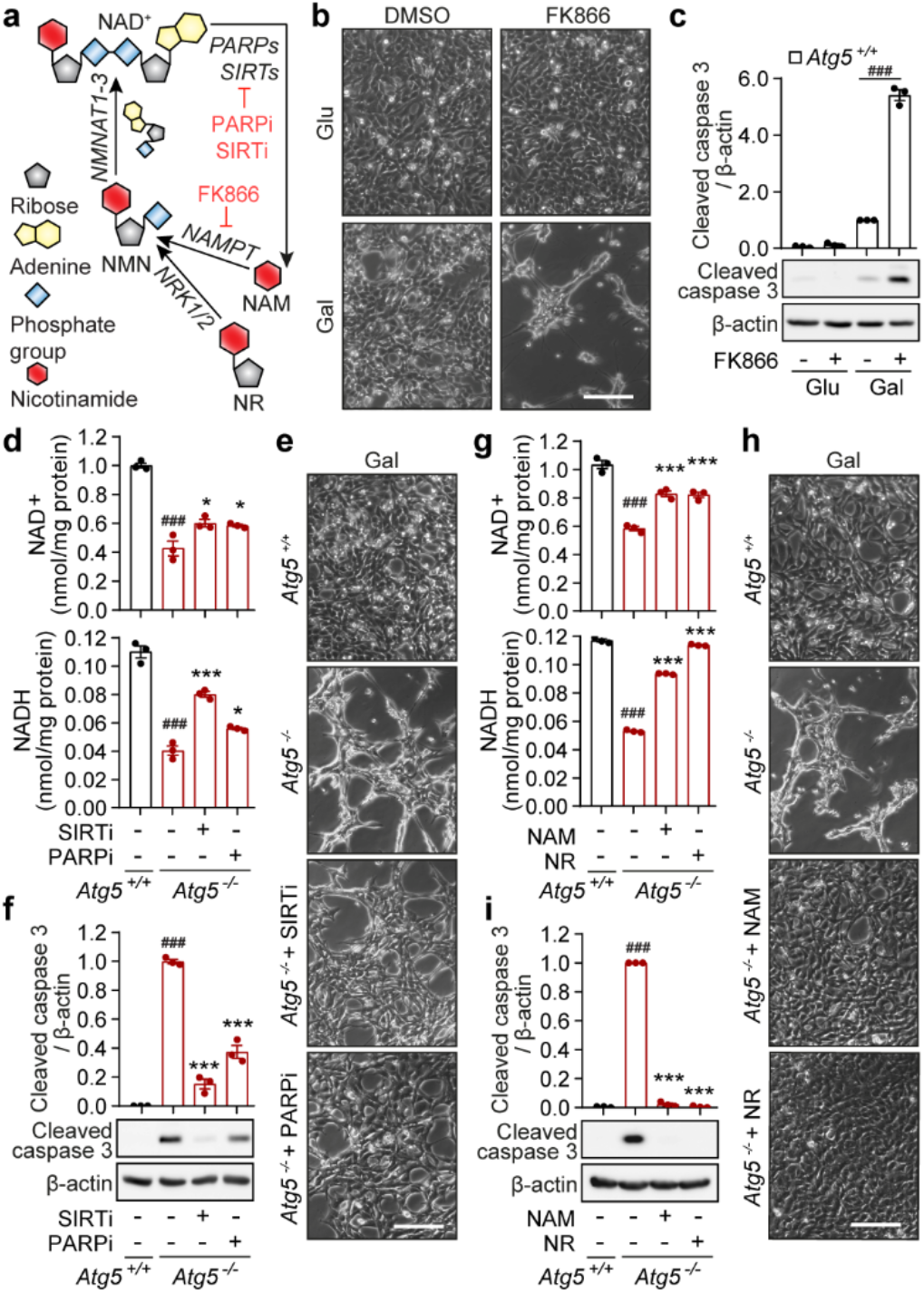
Boosting cellular NAD(H) levels rescues cellular viability. **a**, Graphical representation of the NAD^+^ salvage pathway. Enzyme inhibitors are highlighted in red. **b**, **e**, **h**, Phase-contrast images of *Atg5*^+/+^ MEFs at 48 h after treatment with 10 nM FK866 or solvent (DMSO) cultured in glucose (glu) and gal media (**b**), of *Atg5*^+/+^ and *Atg5*^−/−^ MEFs cultured in gal medium supplemented with NAD^+^ precursors, 5 mM NAM and 5 mM NR (**e**), or of *Atg5*^+/+^ and *Atg5*^−/−^ MEFs cultured in gal medium treated with 20 μM sirtinol (SIRTi), 10 μM olaparib (PARPi) or solvent (DMSO) (**h**). **c**, **f**, **i**, Immunoblot analyses using cleaved caspase 3 and β-actin antibodies in the same conditions as (**b**, **e**, **h**), respectively**. d**, **g**, Measurement of NAD^+^ and NADH levels in *Atg5*^+/+^ and *Atg5*^−/−^ MEFs after 20 h culture in the same conditions as (**e**, **h**), respectively. Data (**c**, **d**, **f**, **g**, **i**) are mean ± s.e.m. *P* values were calculated by unpaired two-tailed Student’s *t*-test (**c**, **d**, **f**, **g**, **i**) on three independent experiments. ^###^*P*< 0.001 (relative to *Atg5*^+/+^), \**P*<0.05, \*\*\**P*<0.001 (relative to *Atg5*^−/−^). Scale bars, 200 μm (**b**, **e**, **h**).

An alternative way of boosting intracellular NAD(H) levels, is to utilize the native cellular capacity for NAD^+^ synthesis from circulating precursors. Indeed, supplementation of bioavailable NAD^+^ precursors, NAM or nicotinamide riboside (NR), led to the recovery of intracellular NAD^+^ and NADH levels, and completely rescued viability of *Atg5*^−/−^ MEFs (Fig. 2g-i). Importantly, NADH was the only intracellular metabolite that correlated with *Atg5*^−/−^ MEF viability (i.e. it was first found to be significantly depleted in *Atg5*^−/−^ MEF and then rescued by NAD^+^ precursor supplementation; Extended Data Fig. 2c-e). Therefore, we speculated that it may be the loss of NADH that underlies cell death as a result of autophagy deficiency.

Whilst the total intracellular NAD(H) pool can only be depleted by enzyme-assisted cleavage of the oxidised form, NAD^+^, the loss of intracellular NADH levels can be further exacerbated by its oxidation to NAD^+^ by the mitochondrial NADH:ubiquinone oxidoreductase (complex I; CI). We hypothesized that depletion of the mitochondrial NADH pool in respiring mitochondria was instrumental for triggering apoptosis, as multiple observations in our study were consistent with the role of mitochondria in the execution of cell death. First, we found that cell death occurs only when *Atg5*^−/−^ MEFs generate energy via mitochondrial OXPHOS (Fig. 1), and second, NADH depletion in autophagy-deficient cells cultured in galactose medium takes place primarily within mitochondria (Fig. 3a). To test if increasing mitochondrial NADH oxidation exacerbates the cell death phenotype of *Atg5*^−/−^ MEFs, we overexpressed a non-proton pumping alternative NADH oxidase, NDI1^24^. When cultured in galactose medium, overexpression of NDI1 enhanced the apoptotic phenotype of autophagy-deficient cells (Extended Data Fig. 3a, b). In contrast, suppression of OXPHOS by cell culture under hypoxia rescued cell death of *Atg5*^−/−^ MEFs in galactose medium (Extended Data Fig. 3c, d). Additionally, knockdown of subunits within the CI and CIII of the electron transport chain (but not CII, which is not involved in NADH oxidation) also promoted survival of *Atg5*^−/−^ MEFs (Extended Data Fig. 3e-h). Together, these data suggest that cellular dependency on NADH oxidation through CI-CIII-dependent mitochondrial respiration contributes to cell death as a result of autophagy deficiency.

**Fig. 3:**
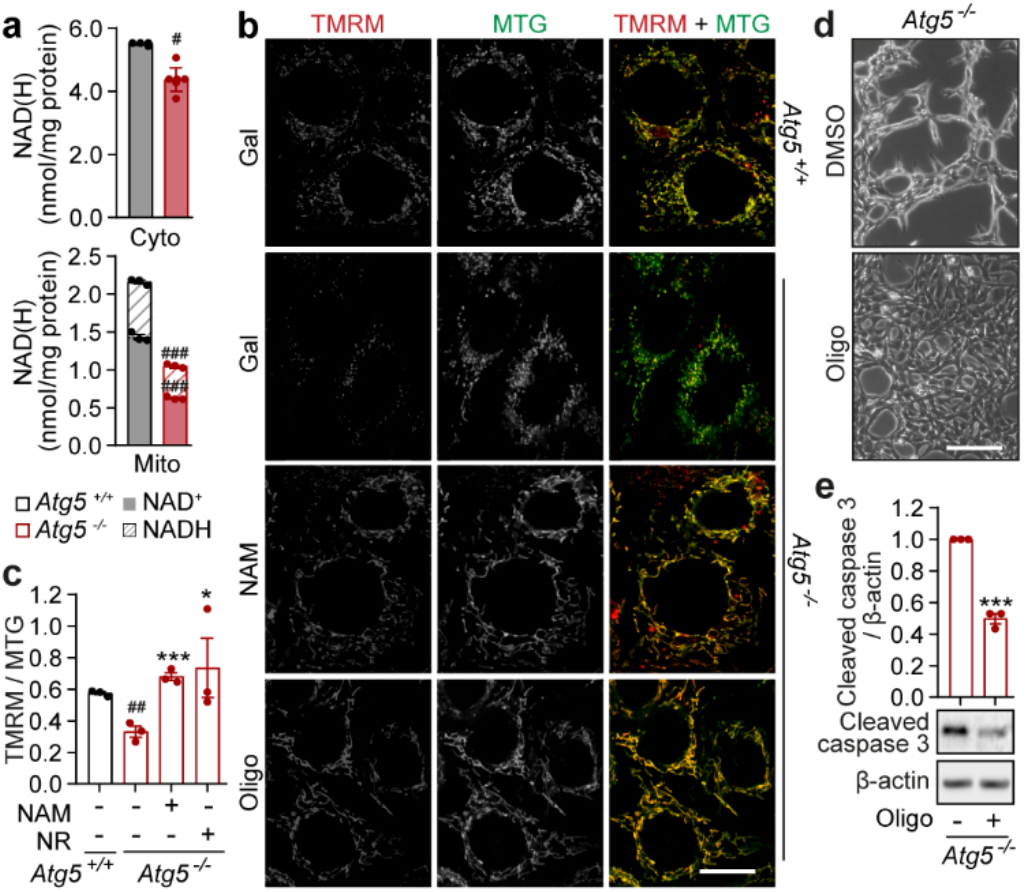
Loss of NAD(H) results in mitochondrial depolarisation and cell death of *Atg5*^−/−^ MEFs. **a**, Measurement of NAD^+^ and NADH levels in cytoplasmic (cyto) and mitochondrial (mito) fractions of *Atg5*^+/+^ and *Atg5*^−/−^ MEFs after 20 h culture in gal medium **b**, Confocal immunofluorescence images of live cells supplemented with 5 mM NAM or 1 nM oligomycin (Oligo) co-stained with TMRM and MTG by direct addition of both dyes to gal medium. **c**, ΔΨm was quantified as a ratio of TMRM to MTG. **d**, Phase-contrast images of *Atg5*^−/−^ MEFs cultured in gal medium supplemented with 1 nM Oligo or solvent (DMSO). **e**, Immunoblot analysis using cleaved caspase 3 and β-actin antibodies in the same conditions as (**d**). Data (**a**, **c**, **e**) are mean ± s.e.m. *P* values were calculated by unpaired two-tailed Student’s *t*-test (**a**, **c**, **e**) on three independent experiments. ^#^*P*<0.05, ^##^*P* < 0.01, ^###^*P*< 0.001 (relative to *Atg5*^+/+^), \**P*<0.05, \*\*\**P*<0.001 (relative to *Atg5*^−/−^). Scale bar, 20 μm (**b**) and 200 μm (**d**).

We next asked how the depletion of NADH in mitochondria could lead to cell death. Oxidation of NADH is used to generate mitochondrial membrane potential (ΔΨm) across the inner mitochondrial membrane that is subsequently consumed by mitochondrial ATP synthase to power ATP production. We hypothesized that depletion of mitochondrial NADH would lead to insufficient ΔΨm generation, which is known to trigger mitochondrial recycling via autophagy^25^. However, absence of autophagy would lead to persistent mitochondrial depolarization and apoptosis^26^. Confirming our hypothesis, we found that ΔΨm was significantly decreased in respiring autophagy-deficient cells and boosting NAD(H) levels, by supplementing media with NAM, rescued membrane depolarization in *Atg5*^−/−^ MEFs (Fig. 3b, c). Importantly, preventing dissipation of ΔΨm, by suppressing ATP synthase activity (using low doses of oligomycin), was sufficient to decrease levels of cell death (Fig. 3d, e). Considering all the evidence, we conclude that depletion of NADH within mitochondria of autophagy-deficient cells leads to the loss of ΔΨm, and triggers caspase activation and apoptosis.

Autophagy is required for the survival of eukaryotic organisms from yeast to man^27^. We investigated whether the key role of autophagy in the maintenance of intracellular NAD(H) pools we discovered in MEFs is evolutionarily conserved and if it underlies the importance of autophagy for organismal survival. In agreement with others^28^, we found that in a yeast model, *Saccharomyces cerevisiae,* autophagy is upregulated in response to nitrogen deprivation and is required for survival in starvation conditions (Fig. 4a and Extended Data Fig. 4a). Loss of viability of *atg5* knockout *S. cerevisiae* was preceded by a striking depletion of both NAD^+^ and NADH and supplementation with NAM partially rescued both, NAD(H) levels and cell survival (Fig. 4a, b). Similarly, knockdown of *ATG5* in the fruit fly *Drosophila melanogaster* resulted in a shortened median lifespan and NAD(H) depletion, which was partially rescued by supplementation with NAM (Fig. 4c, d). Importantly, NAM-mediated protection from loss of viability in either model did not occur as a result of autophagy reconstitution (Extended Data Fig. 4a, b). This finding is consistent with the protective effects of NAD(H) boosting strategies, downstream of autophagy dysfunction, that we observed in our cell-based models. To test the relevance of our findings from genetic knockout/knockdown systems to human disease presenting with an autophagy impairment, we analysed primary fibroblasts derived from patients suffering with a form of lysosomal storage disease, the Niemann-Pick type C1 disorder, caused by mutations in a cholesterol transporter NPC1^29^ (Extended Data Fig. 4c-f). Although primary fibroblasts from patients carrying *NPC1* mutations did not display a spontaneous cell death phenotype in galactose media (potentially due to only a partial loss of autophagy function), they did display decreased NAD(H) levels and an increased sensitivity to exogenous stress in these conditions (Fig. 4e, f). As in the other cell and animal models tested above, both the NAD(H) deficit and the increased sensitivity to stress could be rescued by supplementation with NAM, without alleviating the underlying autophagy defect (Fig. 4e, f and Extended Data Fig. 4g). Based on our findings in model organisms and patient-derived fibroblasts, we conclude that loss of NAD(H) homeostasis is a key mechanism by which autophagy dysfunction leads to the loss of cellular and organismal viability. As such, our data reveal a novel and evolutionarily conserved role of autophagy in the maintenance of the cellular NAD(H) pool.

**Fig. 4:**
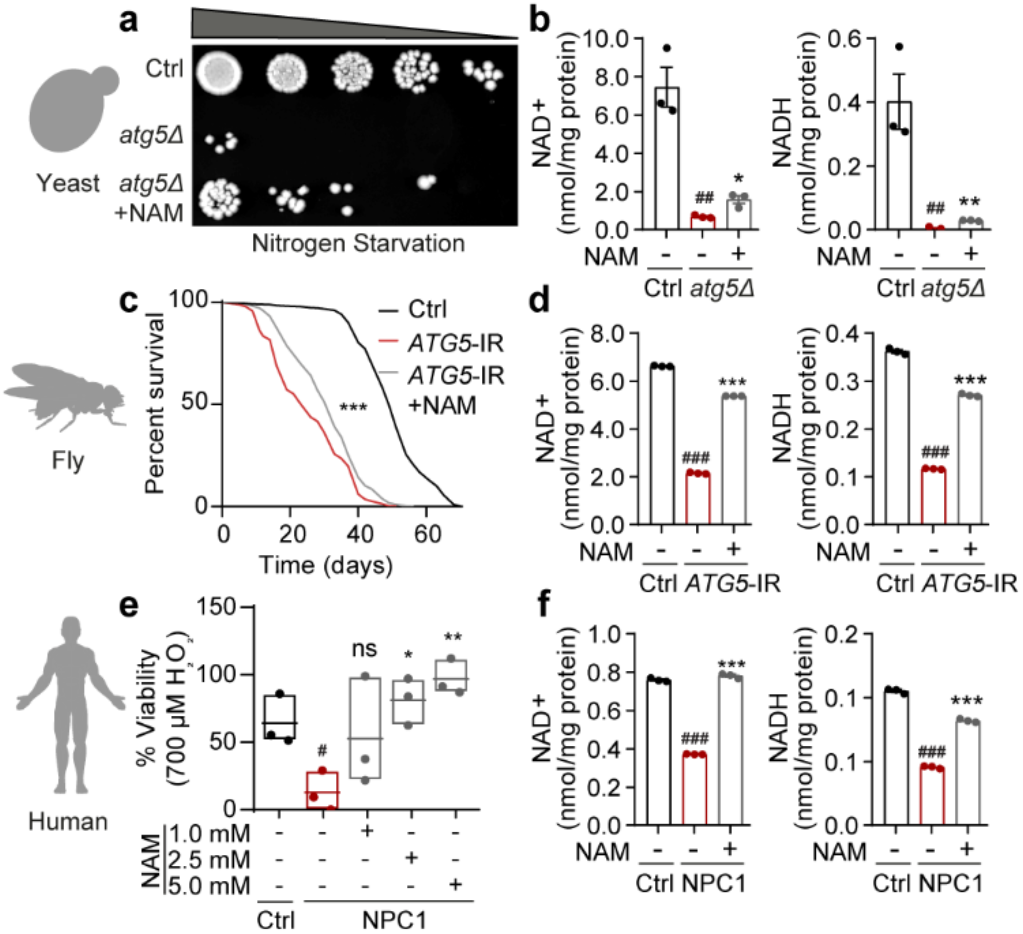
Autophagy has an evolutionarily conserved role in the maintenance of NAD(H) pools and cellular/organismal viability. **a**, Spot-testing of ctrl and *atg5Δ* yeast strains supplemented with 5 mM NAM or solvent (H_2_O) on nutrient-rich agar media following 5 days of nitrogen starvation. Image is a representative of *n*=4 experiments. **b**, Measurement of NAD^+^ and NADH levels in yeast cells at day 5 of nitrogen starvation. **c**, Combined survival data of ctrl (daGAL4>ATTP2) and *ATG5*-IR (daGAL4>*ATG5*-IR) flies fed on standard media with and without NAM cultured at 25 ^o^C (*n*=100 flies per group). **d**, Measurement of NAD^+^ and NADH levels in flies collected at day 10 (*n*=20 flies). **e**, MTT-assay based measure of cell viability in human fibroblasts, isolated from healthy volunteers (ctrl) and NPC1 (NPC1) patients, sub-cultured in gal medium supplemented with increasing doses of NAM and challenged with 700 μM H_2_O_2_ for 2 h (n=3 cell lines per group). **f**, Measurement of NAD^+^ and NADH levels in human cells in the same conditions as (**e**) prior to challenge with H_2_O_2_. Details of ctrl and NPC1 patient fibroblast cell lines are listed in Supplementary Table 1. Data are mean ± s.e.m (**b**, **d**, **f**) or box plot (line at median) (**e**). *P* values were calculated by unpaired two-tailed Student’s *t*-test (**b**, **d**, **e**, **f**) and the survival curve comparison log-rank Mantel Cox test on two (**e**) or three independent experiments (**b**, **d**, **f**). ^#^*P*<0.05, ^##^*P*< 0.01, ^###^*P*< 0.001 (relative to ctrl), \**P*<0.05, \*\**P*<0.01, \*\*\**P*<0.001 (relative to model of autophagy deficiency (*atg5Δ*/*ATG5*-IR/NPC1)).

Autophagy is a catabolic pathway crucial for cellular function and survival. Stimulated by a variety of toxic and nutritional stresses, autophagy serves to sequester and degrade damaged cytoplasmic components and recycle their building blocks, including amino acids, lipids and nucleotides, to sustain cellular growth and viability. Thus predictably, decreased levels of nucleotides and loss of cell viability were previously observed in autophagy-deficient cells and tissues^3–5,18^. However, the contribution, or lack thereof, of individual nucleotides and the underlying mechanisms by which their depletion leads to cell death and tissue dysfunction remained unknown. In agreement with others, we have observed a decrease in a range of free nucleotides, which may be caused by defective recycling of nucleic acids in autophagy-deficient cells^18^. Importantly, we demonstrate that it is specifically depletion of the NAD(H) pool that is critical to loss of cell viability, due to mitochondrial depolarisation downstream of autophagy dysfunction.

Ageing and age-related diseases have long been associated with a decline in both autophagy and NAD^+^ levels^29, 30^. Nutritional or pharmacological activation of autophagy is currently a subject of intense research and development of small molecule modulators^7^. However, due to the varied nature of autophagy dysfunction in genetic and age-related sporadic diseases, development of a universal modulator remains unlikely^31^. In contrast, boosting the levels of NAD^+^ by precursor supplementation in animal models was found to have a positive impact on age-related phenotypes, which is at least in part, mediated by upregulation of autophagy^32^. Crucially, our data show that autophagy is, in turn, required for NAD(H) maintenance and that increasing NAD(H) levels protects cells by preventing the loss of ΔΨm even in the absence of functional autophagy. As such, our investigation defines a novel mechanism linking autophagy, NAD(H) metabolism and ageing. Finally, our study establishes several points of intervention that could be targeted therapeutically, in order to alleviate cellular pathology in a range of diseases associated with autophagy dysfunction.

## End Notes

We are grateful to E. Bennett and Newcastle Bioimaging Unit for technical assistance. This work was funded by BBSRC (V.I.K., A.S., M.P.C), MRC CAV (V.I.K.), Wellcome Trust (S.S.) and LifeArc (S.S.). O.D.K.M. is funded by a Cancer Research UK Career Development Fellowship (C53309/A19702). T.K. was supported by Japan Society for the Promotion of Science fellowship. R.S is a Sir Henry Wellcome Postdoctoral Fellow funded by Wellcome (204715/Z/16/Z). ET and ST were funded by the National Institutes of Health (NIA RF1AG55549 and NINDS R01NS107265 grants to ET).

## Author Contributions

L.S., E.G.O., B.C., T.K., E.S., Y.R-R., G.K., N.K. and F.U. performed cell biology experiments; F.S. and R.S. performed fly experiments; D.S. performed yeast experiments; A.A. generated bioinformatics data; G.N. supervised and performed bioimaging experiments; T.Z. and O.D.K.M. performed mass spectrometry; L.S., E.G.O., S.T., C.C.B, R.T, R.J.I., J.E.O., E.T, M.I., S.S., M.L., M.P.C., O.D.K.M., S.S., A.S., and V.I.K. designed and supervised elements of the study; S.S., A.S., and V.I.K. supervised the entire project and wrote the paper with help from all authors.

## Experimental Procedures

### Culture of mammalian cells and human fibroblasts

*Atg5*^+/+^, *Atg5*^−/−^ MEFs (gift from Noboru Mizushima^33^), *Npc1*^+/+^ and *Npc1*^−/−^ MEFs (gift from Peter Lobel^34^) were maintained in DMEM (Sigma) supplemented with 10% foetal bovine serum (FBS) (Sigma), 100 U/ml penicillin/streptomycin (Sigma) and 2 mM L-glutamine (Sigma) at 37 °C, and 5% CO_2_ in a humidified incubator. HEK293T cells, purchased from Life Technologies (R700-07), were cultured as above in medium supplemented with 1x MEM non-essential amino acids (Sigma). Control young female human fibroblasts (10156, 10263 and 10632) (a kind gift from Devin Oglesbee) and NPC1 patient fibroblasts (GM18381, GM18402 and GM18417) obtained from Coriell Cell Repositories were cultured as above, except with 15% FBS. Details of primary human fibroblasts are listed in Supplementary Table 1.

### *Saccharomyces cerevisiae* stocks and viability experiments

S288C (*MATα SUC2 gal2 mal2 mel flo1 flo8-1 hap1 ho bio1 bio6)* WT and *atg5Δ::KanMX* (gift from Charles Boone^35^) were maintained in synthetic complete medium (SC medium: 0.13% drop-out CSM powder (Formedium), 0.17% yeast nitrogen base (Formedium), 2% glucose (Formedium), 0.5% ammonium sulphate (Sigma) to mid/late log phase (OD_600_ 0.8 – 1) at 30 ⁰C. For assessment of viability, both strains were washed in dH_2_O and switched to a nitrogen starvation medium (SD-N medium: 0.17% yeast nitrogen base), or SD-N medium supplemented with 5 mM nicotinamide (Sigma). Strains in all conditions were grown for 5 days and either collected by snap freezing in liquid nitrogen for NAD(H) measurement assay or processed for a spot-test assay. The individual strain concentrations were equalised by OD_600_ and subsequently spotted on to YEPD plates (YEPD broth (1% yeast extract, 2% peptone, 2% glucose) Formedium), 2% agar) in a 5-fold serial dilution and left to grow at 30 ⁰C for 48 h prior to imaging in G-box transilluminator (Syngene).

### *Drosophila melanogaster* stocks and lifespan experiments

Daughterless-GAL4, Atg5-IR (BL34899) and the control Attp2 (BL36303) were obtained from the Bloomington Drosophila Stock Center (BDSC). Flies were crossed and cultured on standard media (1% agar (SLS), 1.5% sucrose (VWR), 3% glucose (VWR), 3.5% active dried yeast (SLS), 1.5% white maize meal (Asda), 1% wheatgerm (MP biomedicals), 1% soybean flour (Santa Cruz Biotechnology), 3% treacle (Bidfood), 0.5% propionic acid (VWR), 0.1% Nipagin (Sigma) or on standard media supplemented with 5 mM of nicotinamide (Sigma). Eclosed male flies were collected using CO_2_ anaesthesia and maintained at a density of 20 flies per vial at 25°C. Flies were transferred to fresh vials every 2-3 days. A minimum of 100 flies per genotype were used for the lifespan experiments and repeated three times. The number of dead flies was recorded every 2-3 days, and the median lifespan was calculated for each experiment. The data were analysed by using the survival curve comparison log-rank Mantel Cox test in GraphPad Prism 8 software.

### Generation of transient and stable cell lines

Re-introduction of the *Atg5* gene into *Atg5*^−/−^ MEFs, was achieved by packaging retroviruses in the HEK293FT (293FT) cell line. 293FT cells were seeded in a 10 cm dish (6.0×10^6^ cells/10ml/dish) in antibiotic-free glucose culture medium. Next day, cells were transfected with plasmids containing the packaging psPAX2 (Addgene, 12260, from Didier Trono) and envelope pCMV-VSV-G (Addgene, 8454, from Bob Weinberg^36^) genes, and the pMXS-IP-eGFP (Addgene, 38192, from Noboru Mizushima^37^) or pMXs-IP-eGFP-mAtg5 (Addgene, 38196, from Noboru Mizushima^37^) constructs with Lipofectamine® 2000 transfection reagent (Invitrogen). Following overnight transfection, the medium was replaced with fresh antibiotic-free medium that was collected after 24 h. Virus containing medium was filtered through at 0.45 μm pore-size filter and overlaid on 70% confluent *Atg5*^−/−^ MEFs for 24 h in the presence of 10 μg/ml polybrene (Sigma). Cells stably expressing the *Atg5* gene were optimised for protein expression via 2 μg/ml puromycin (Santa Cruz Biotechnology) selection for 7 days.

Introduction of the *NDI1* gene into *Atg5*^−/−^ MEFs was achieved by transient cell transfection. *Atg5*^−/−^ MEFs were seeded in a 6-well plate, cultured for 24 h and transfected with either pWPI-eGFP (Addgene, 12254, from Didier Trono) or pWPI-eGFP-NDI1^38^ constructs with the with the Lipofectamine® 2000 transfection reagent (Invitrogen) with 1.6 μg plasmid DNA according to manufacturer instruction 48 h prior to passaging and galactose culture.

### Generation of knockout lines using CRISPR/Cas9 gene editing

*Atg5*^−/−^, *Atg7*^−/−^ and *Rb1cc1*^−/−^ MEFs were generated using the clustered regularly interspaced short palindromic repeats (CRISPR)/Cas9 system. Ensembl, Aceview and CHOPCHOP databases were utilised to design CRISPR guide RNAs (gRNAs) to target exons present in all splicing variants of the targeted gene (Supplementary Table 2). The gRNA oligomer was then annealed and ligated into Bbsl (Fisher Scientific) linearized pSpCas9(BB)-2A-GFP gRNA vector^39^. WT MEF cell line seeded into a 6-well plate was then used for transfection with DNA ligation products. Cells were allowed to grow for 24 h post seeding before transfection with Lipofectamine® 2000 (Invitrogen) with 1.6 μg plasmid DNA according to manufacturer’s instructions. GFP-positive cells were sorted by FACS into 96-well plates and expanded into colonies prior to screening for autophagy impairment by immunoblotting.

### Transient knockdown of genes using siRNA

ON-TARGETplus SMARTpool siRNA against mouse *Ndufs3* (L-047009-01-0005), *Sdha* (L-046818-01-0005), *Uqcrfs1* (L-057582-01-0005) were purchased from Dharmacon. Final siRNA concentration of 100 nM was used for silencing, and transfections were performed with Lipofectamine® 2000 (Invitrogen) as per company instructions. Cells were analysed 72 h post-transfection

### Galactose Medium Culture and Supplementation

To induce mitochondrial respiration, cells seeded in a 6-well format (0.3×10^6^ cells/2ml/well) were switched to a galactose medium (glucose-free DMEM (Gibco) supplemented with 10 mM D-galactose (Sigma), 10 mM HEPES (Sigma), 1 mM sodium pyruvate (Sigma), 4 mM L-glutamine (Sigma), 100 U/ml penicillin/streptomycin (Sigma) and 10% FBS (Sigma)) 24 h post-seeding. Galactose medium was supplemented with various compounds and inhibitors: 20 μM Z-VAD-fmk (Enzo), 10 nM FK866 (Sigma), 5 mM NAM (Sigma), 5 mM NR (ChromaDex), 10 μM olaparib (Cambridge Biosciences), 20 μM sirtinol (Cambridge Biosciences), 400nM bafilomycin A_1_ (Enzo Life Sciences), 10 nM oligomycin (Merck), or 400-700 μM H_2_O_2_ (Sigma). All compound supplements were added at 0 h, except Z-VAD-fmk which was supplemented at 20 h and H_2_O_2,_ which was added to primary human fibroblasts after a 4-passage subculture in galactose medium. For hypoxia experiments, cells were incubated at 1% O_2_ in an *in vivo* 400 hypoxia work station (Ruskin, UK). Cells were lysed for protein extracts in the chamber to avoid re-oxygenation.

### MS-based metabolomics

Metabolite extraction for liquid-chromatography-mass spectroscopy (LC-MS) was performed on MEFs following a 20 h incubation in galactose medium. Cells were washed once with cold PBS (New England Biolabs) and lysed at a concentration of 2×10^6^ cells/ml in a metabolite extraction buffer (50% methanol (Fisher Scientific), 30% acetonitrile (Sigma), 20% dH_2_O). Samples were vortexed for 45 s, centrifuged at 16,100 g and supernatants subjected to LC-MS as follows, using a three-point calibration curve with universally labelled carbon-13/nitrogen-15 amino acids for quantification

Prepared samples were analysed on a LC-MS platform consisting of an Accela 600 LC system and an Exactive mass spectrometer (Thermo Scientific). A Sequant ZIC-pHILIC column (4.6mm x 150mm, 5μm) (Merck) was used to separate the metabolites with the mobile phase mixed by A=20mM ammonium carbonate in water and B=acetonitrile. A gradient program starting at 20% of A and linearly increasing to 80% at 30 min was used followed by washing (92% of A for 5 mins) and re-equilibration (20% of A for 10min) steps. The total run time of the method was 45 min. The LC stream was desolvated and ionised in the HESI probe. The Exactive mass spectrometer was operated in full scan mode over a mass range of 70– 1,200 m/z at a resolution of 50,000 with polarity switching. The LC-MS raw data was converted into mzML files by using ProteoWizard and imported to MZMine 2.10 for peak extraction and sample alignment. A house-made database integrating KEGG, HMDB and LIPID MAPS was used for the assignment of LCMS signals by searching the accurate mass and the metabolites used in the manuscript were confirmed by running their commercial standards. Finally, peak areas were used for comparative quantification.

### Separation of cytoplasmic and mitochondrial fractions in mammalian cells

Mitochondria were isolated from a total of ~60 million (30×6-well) cells by manual cell homogenization in a specialised buffer (20 mM HEPES (pH 7.6) (Sigma), 220 mM D-mannitol (Sigma), 70 mM sucrose (Sigma), 1 mM EDTA (Sigma), 0.5 mM phenylmethylsulfonyl fluoride (Sigma) and 2 mM dithiothreitol (DTT) (Supelco)). Cell homogenates were centrifuged thrice at 800 *g* at 4 °C for 5 min to pellet cellular nuclei and membrane debris. Cytoplasmic and mitochondrial fractions were separated by centrifugation at 16 000 *g* at 4 °C for 10 min and immediately processed for an NAD^+^ and NADH measurement.

### NAD+ and NADH measurements

Measurements of NAD^+^ and NADH in mammalian whole cell lysates, mitochondrial lysates, and in whole yeast strains and flies were performed as described in a published protocol^*40*^. In brief, NAD^+^ or NADH were extracted with an acidic solution (10% (mitochondria, mammalian cell, yeast) or 20% (fly) trichloroacetic acid (TCA) (Sigma) or basic solution (0.5 M sodium hydroxide (NaOH, Sigma), 5 mM EDTA(Sigma)) respectively. NAD^+^ and NADH pools from cellular cytoplasmic fractions were extracted by addition of 5x concentrated stocks of TCA and NaOH/EDTA solutions into the cytoplasmic supernatant. Samples were adjusted to pH 8 with 1 M Tris (Sigma). NAD^+^ and NADH levels were determined by fluorescence intensity of resorufin produced by an enzymatic cycling reaction using resazurin (Sigma), riboflavin 5’-monophosphate (Sigma), alcohol dehydrogenase (Sigma) and diaphorase (Sigma). Fluorescence intensity was monitoredevery minute for a total 60 min using a microplate reader (FLUOstar Omega, BMG Labtech). NAD^+^ and NADH levels were determined by a β-NAD (Sigma) standard curve and adjusted to protein concentration determined by the DC protein assay (BioRad).

### Mitochondrial ΔΨ measurements

Cells were grown in 96-well glass bottom dishes (Greiner Bio-One) (0.8×10^4^ /100 μl/well, 24 h). Following culture in galactose medium for 40 h, cells were co-stained with 16.7 nM tetramethylrhodamine methyl ester (TMRM; Invitrogen) and 100 nM Mitotracker Green (MTG; Invitrogen). A 10x stock of each compound was prepared in conditioned galactose medium (24 h culture on *Atg5*^−/−^ MEFs, collected and filtered through a 0.22 μm pore-size filter) and added directly to culture wells. Following a 30 min incubation in the dark at 37 °C, TMRM- and MTG-containing medium was replaced by dye-free conditioned galactose medium. Live cell imaging was performed in a maintained atmosphere of 37 °C and 5% CO_2_ using an LSM700 microscope (Zeiss) with a C-Apochromat 40x/1.20 water immersion lens, capturing images line sequentially.

### MTT assay following H_2_O_2_ treatment

Cell viability upon H_2_O_2_ treatment was measured indirectly by a high-throughput MTT assay. Human fibroblasts were seeded into a 96-well plate (0.1×10^4^ cells/100 μl/well) following a 4-passage subculture in a galactose medium. Cell viability was challenged by 700 μM H_2_O_2_ addition 48 h after seeding. After a 2 h incubation following the H_2_O_2_ addition, 2.5 μg/ml thiazolyl blue tetrazolium bromide (MTT, Sigma) was added to each well and cells were incubated in the dark at 37 °C for two hours.

Formazan crystals were then solubilised by addition of HCl (Sigma) in isopropanol (Sigma) at a final concentration of 25 mM followed by a 10 min shaking incubation at RT. The absorbance was read at 570 nm on a multi-well plate reader (BMG Labtech). Reduced formazan quantitation values were first corrected by protein levels measured from a duplicate plate and then normalised to an internal control of non-H_2_O_2_ treated cells (100% viable).

### Assessment of autophagy

#### Human fibroblasts

Autophagic flux in primary human fibroblasts was first assessed by immunoblot and immunofluorescence analyses upon culture in glucose medium. Cells were seeded for immunoblot (0.12×10^6^ cells/2 ml/6-well) or immunofluorescence (0.12×10^6^ cells/2 ml/6-well with 2×13 mm coverslips) analyses. Samples for immunofluorescence analysis were processed after 48 h of culture. For immunoblot-based analysis of autophagy flux, bafilomycin A1 (bafA_1_) was added directly to cell culture media at a final concentration of 100 nM after 44 h culture. Cells were processed for immunoblotting at 48 h following a 4 h incubation with bafA_1_. For immunoblot assessment of LC3B lipidation levels in cells cultured in galactose medium, cells were collected following a 4-passage subculture in galactose-medium alone or supplemented with 5 mM NAM every 2-3 days.

#### S. cerevisiae

Autophagic activity was assessed upon transformation of WT and *atg5Δ::KanMX* strains with a GFP-ATG8(416)/GFP-AUT7(416) plasmid (Addgene, 49425, from Daniel Klionsky^41^). Transfected strains were switched to SD-N media and collected prior to the switch (0 h) and at 2 h, 4 h and 18 h time-points following the switch. Samples equivalent to 5 ml at OD_600_ 1, were taken at the indicated times for protein extraction and immunoblot analysis.

#### D. melanogaster

Levels of Ref(2)P as a proxy for autophagy-mediated degradation of intracellular substrate were probed by immunoblotting 10 d whole fly lysates.

### Cell death assays

Adherent and floating cells were collected and processed by protein extraction and immunoblot analysis at 24 h (*Atg5*^−/−^), 48 h (*Atg5*^+/+^), 72 h (*Npc1*^+/+^ and *Npc1*^−/−^) and 110 h (all CRISPR-Cas9 generated cell lines) after media switch. Representative phase-contrast images were obtained on an inverted DM-IL Leica microscope equipped with an Invenio 3SII digital camera (3.0 Mpix Colour CMOS; Indigo Scientific).

### Immunoblot analysis

#### Mammalian cells

Immunoblotting on cells was performed as described previously^42^. In brief, cells were lysed on ice in RIPA buffer (Sigma) supplemented with 1x Halt^TM^ protease and phosphatase inhibitor cocktail (Thermo Scientific). Protein concentration of lysates was measured using DC protein assay (Bio-Rad Laboratories), and equal amounts of protein (20–40 μg) were subjected to SDS-PAGE.

#### S. cerevisiae

Yeast sample preparation for immunoblotting based on a TCA protein extraction protocol. 10ml cultures were grown in the appropriate medium to an OD_600_ of 0.8. Cells were pelleted by centrifugation and washed with 20% TCA (Sigma). All of the following purification steps were performed on ice with pre-chilled solutions. Cell pellets were re-suspended in 100 μl of 20% TCA and subjected to glass bead lysis. The supernatant was collected, 200 μl of 5% TCA was added, and the precipitated proteins were pelleted by centrifugation. Protein pellets were solubilised in 30 μl of 2 M Tris pH 8.0 (Sigma) / 70μl 3X SDS-PAGE loading buffer (60 mM Tris pH 6.8 (Sigma), 2% SDS (Bio-Rad), 10% glycerol (Sigma), 100mM DTT (Sigma), 0.2% bromophenol blue (Sigma)) and boiled at 95 °C for 5 min. Insoluble material was removed by centrifugation and the supernatant was subjected to SDS-PAGE analysis.

#### D. melanogaster

Fly sample preparation and immunoblotting were performed as described previously^42^. Briefly, 20 flies per group were homogenized on ice in a specialised buffer (1.5% Triton X-100 (Promega), complete mini EDTA-free proteinase inhibitor (Sigma), PBS 1X). Protein concentration in the supernatant was measured using Bradford Reagent (Sigma), and equal amounts of protein were resolved by SDS-PAGE.

Membranes were first blocked in 5% milk (Marvel) in PBS-1x Tween^®^ 20 (Sigma) for 1 h at RT and incubated with primary antibodies overnight at 4 °C on a shaker platform (for a full list see Supplementary Table 3). Secondary antibodies conjugated to horseradish peroxidase (HRP) were used at 1:5000 dilution for 1 h at RT. In cell and fly samples, clarity western ECL substrate (Bio-Rad Laboratories) was used to visualise chemiluminescence on LAS4000 (Fujifilm). The chemiluminescent signal in yeast samples was generated by the SuperSignal West Pico Plus chemiluminescent substrate (Thermo Scientific) and detected on a G-box transilluminator (Syngene).

### Immunofluorescence

Immunofluorescence analysis was performed on primary human fibroblasts as described previously^42^. In brief, cells were washed once with 1x PBS (New England Biolabs) and fixed and permeabilised in 100% pre-chilled methanol (Fisher Scientific) for 5 min at −20 °C. Cells were then incubated with blocking buffer (5% goat serum (Sigma) in 1x PBS) for 1 h at RT and incubated with an LC3B antibody (1:200; Cell Signalling Technology, 3868) overnight at 4 °C. Cells were washed and incubated with Alexa Fluor 488 goat anti-rabbit (H+L) antibody (1:1,000; Thermo Fisher Scientific; A-11008) for 1 h at RT. Coverslips were mounted on slides with ProLong^TM^ Gold antifade reagent with DAPI (Fisher Scientific). Fluorescence images were captured on an Axio observer Z1 microscope (Zeiss), with a Plan-Apochromat 20x/0.8 M27 air immersion objective, equipped with an Axiocam 503 camera (Zeiss).

### Quantification and statistical analysis

Output from MS-based metabolomics was subjected to statistical analysis by MetaboAnalyst 4.0. We first performed a multivariate statistical principal component analysis (PCA). The variables were normalised by auto-scaling (mean-centered and divided by SD of each variable) by the MetaboAnalyst platform and then subjected to PCA analysis. Furthermore, a univariate statistical test coupled with *Atg5*^−/−^ / *Atg5*^+/+^ fold change of each metabolite were plotted on a volcano plot. Statistical significance was determined using the Student’s *t-*test with *P* value corrected with false discovery rate (FDR) method of Benjamini and Hochberg which does not assume a consistent standard deviation (SD).

Analysis of images captured by immunofluorescence was performed in ImageJ (version 1.41) (National Institutes of Health (NIH)) by outlining single cells as regions of interest and application of a constant image threshold to determine numbers of LC3 puncta per cell.

Mitochondrial membrane potential image analysis was performed in ImageJ (version 1.41; NIH) by outlining single cells as regions of interest and calculation of a ratio of TMRM to MTG raw integrated density values per cell. Quantification was performed on 30–40 cells per condition in three independent experiments.

Densitometry analyses on immunoblots were performed by ImageJ (version 1.41; NIH) software as described previously^42^. Data of the control condition was normalised to 100% and graphical data denote the mean ± s.e.m. All experiments were carried out in three biological triplicates. Unless indicated otherwise, the *P* values for analyses was determined by Student’s *t* test (two-tailed, unpaired) using Prism 8 software (GraphPad).

## Supplementary tables

**Supplementary Table 1:** Details of primary human fibroblasts

| Human Fibroblast | Catalogue ID | Gender | Age at Sampling | Disease Affected | Mutation |
| --- | --- | --- | --- | --- | --- |
| CTRL (1) | 10263 | ♀ | 21 | No | none |
| CTRL (2) | 10632 | ♀ | 22 | No | none |
| CTRL (3) | 10763 | ♀ | 21 | No | none |
| NPC1 (1) | GM18402 | ♀ | 10 | Yes | [D700N] [F1221fsX] |
| NPC1 (2) | GM18417 | ♀ | 25 | Yes | [I1061T] [I1061T] |
| NPC1 (3) | GM18387 | ♀ | 33 | Yes | [D874V] [Y890X] |

**Supplementary Table 2:**
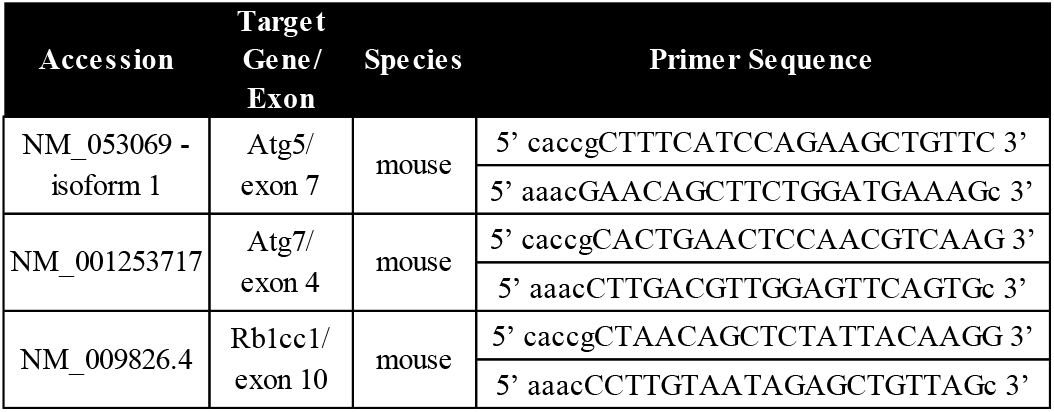
Primers used for CRISPR gRNA generation

**Supplementary Table 3:** List of primary antibodies

| Antigen | Source | Dilution | Catalogue No. |
| --- | --- | --- | --- |
| β-actin | Cell Signalling Technology | 1:10 000 | 4970 |
| Acetylated-Lysine | Cell Signalling Technology | 1:1000 | 9441 |
| Alexa Fluor ® 488 Goat anti-Rabbit IgG *H+L | Invitrogen | 1:500 | A110088 |
| anti-guinea pig HRP | Agilent | 1:5000 | P014102-2 |
| anti-mouse HRP | EMD Biosciences | 1:5000 | 401253 |
| anti-rabbit HRP | EMD Biosciences | 1:5000 | 401393 |
| ATG5-ATG12 | Sigma | 1:1000 | A0856 |
| Cleaved caspase 3 | Cell Signalling Technology | 1:200 | D175 |
| GFP | Roche | 1:2000 | 11814460001 |
| LC3B | Cell Signalling Technology | 1:1000 (WB), 1:200 (IF) | 3868 |
| NDI1 | gift from Takao Yagi's lab (Scripps Institute, CA) |  |  |
| NDUFS3 | Abcam | 1:1000 | ab110246 |
| p62 | Progen Biotechnik | 1:1000 | GP62-C |
| poly(ADP-ribose) | Enzo Life Sciences | 1:1000 | ALX-804-220-R100 |
| Ref(2)p | Abcam | 1:1000 | ab178440 |
| SDHA | New England Biolabs | 1:1000 | 11998 |
| Tubulin | Abcam | 1:5000 | ab179513 |
| UQCRC2 | Abcam | 1:1000 | ab14745 |

## Extended Data Figures

**Extended Data Fig. 1:**
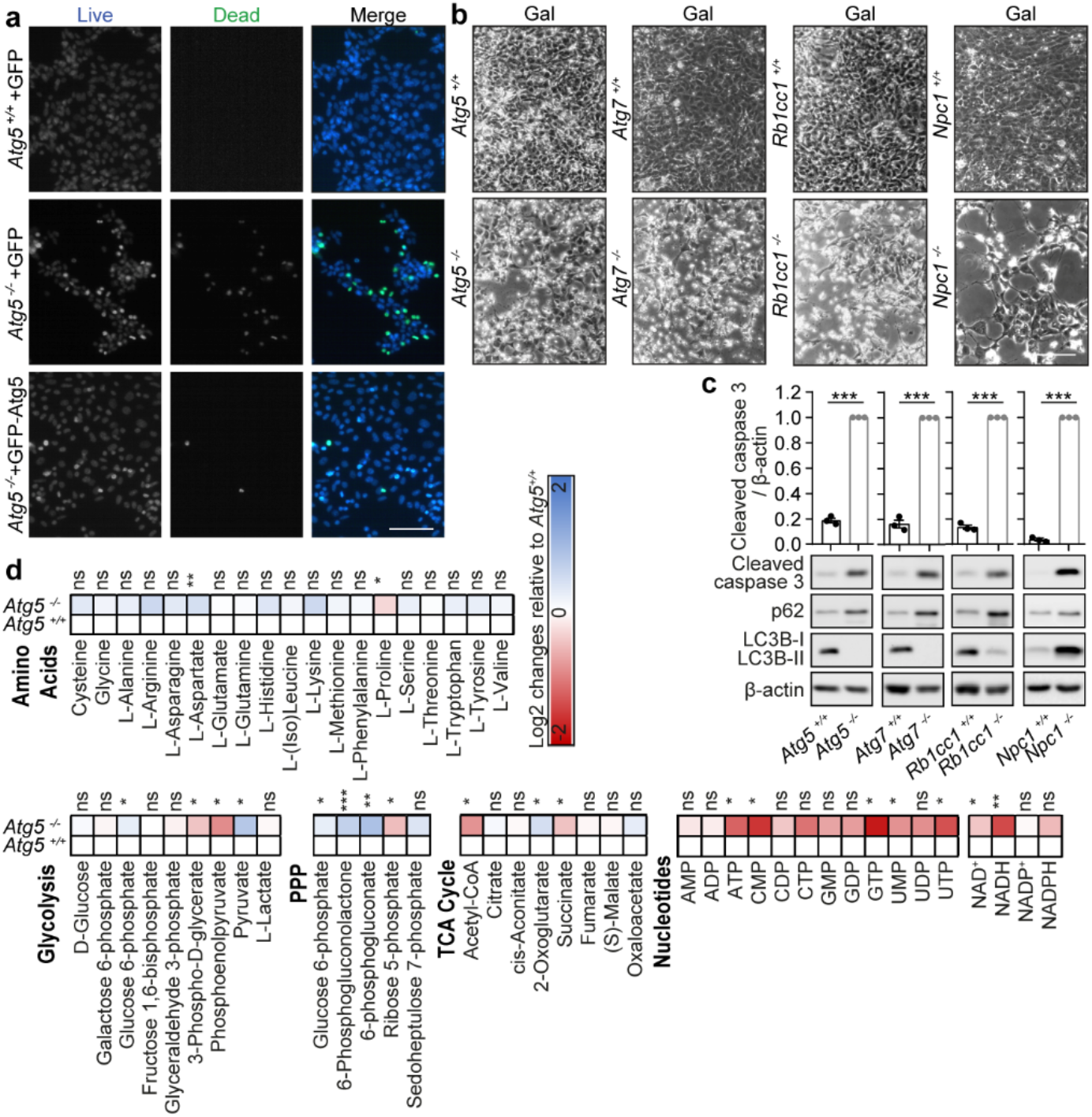
Cell death in galactose medium occurs in multiple autophagy-deficient cell lines. **a**, Staining for cell death with ReadyProbes fluorescent dyes in *Atg5*^+/+^+GFP, *Atg5*^−/−^+GFP and *Atg5*^−/−^+GFP-Atg5 MEFs 24 h after switch to galactose (gal) medium. (*n*=2). **b**, Phase-contrast images of isogenic *Atg5*^+/+^ and *Atg5*^−/−^, *Atg7*^+/+^ and *Atg7*^−/−^, *Rb1cc1*^+/+^ and *Rbcc1*^−/−^ cell lines generated by the CRISPR-Cas9 system; and *Npc1*^+/+^ and *Npc1*^−/−^ MEFs grown in gal media for 110 h (*Atg5, Atg7* and *Rb1cc1* CRISPR-Cas9 generated cell lines) and 72 h (*Npc1* cell lines). **c**, Immunoblot analyses using LC3B, p62, cleaved caspase 3 and β-actin antibodies in the same conditions as (**b**). **d**, Metabolite profiling in *Atg5*^+/+^ and *Atg5*^−/−^ MEFs is depicted as a heatmap of Log2 (fold change (FC)) of *Atg5*^−/−^ to *Atg5*^+/+^ MEFs. Metabolite organisation is based on their association to glucose oxidation pathways of glycolysis, pentose phosphate pathway (PPP) and tricarboxylic acid (TCA) cycle. Amino acids are organised alphabetically. Nucleotide order is first alphabetical and depends on the energy charge. Data (**c**) are mean ± s.e.m. *P* values were calculated by unpaired two-tailed Student’s *t*-test (**c**) and the multiple *t*-test original FDR method of Benjamini and Hochberg (**d**) on three independent experiments. \**P*<0.05, \*\**P*<0.01, \*\*\**P*<0.001, ns (non-significant). Scale bars, 200 μm (**a**, **b**).

**Extended Data Fig. 2:**
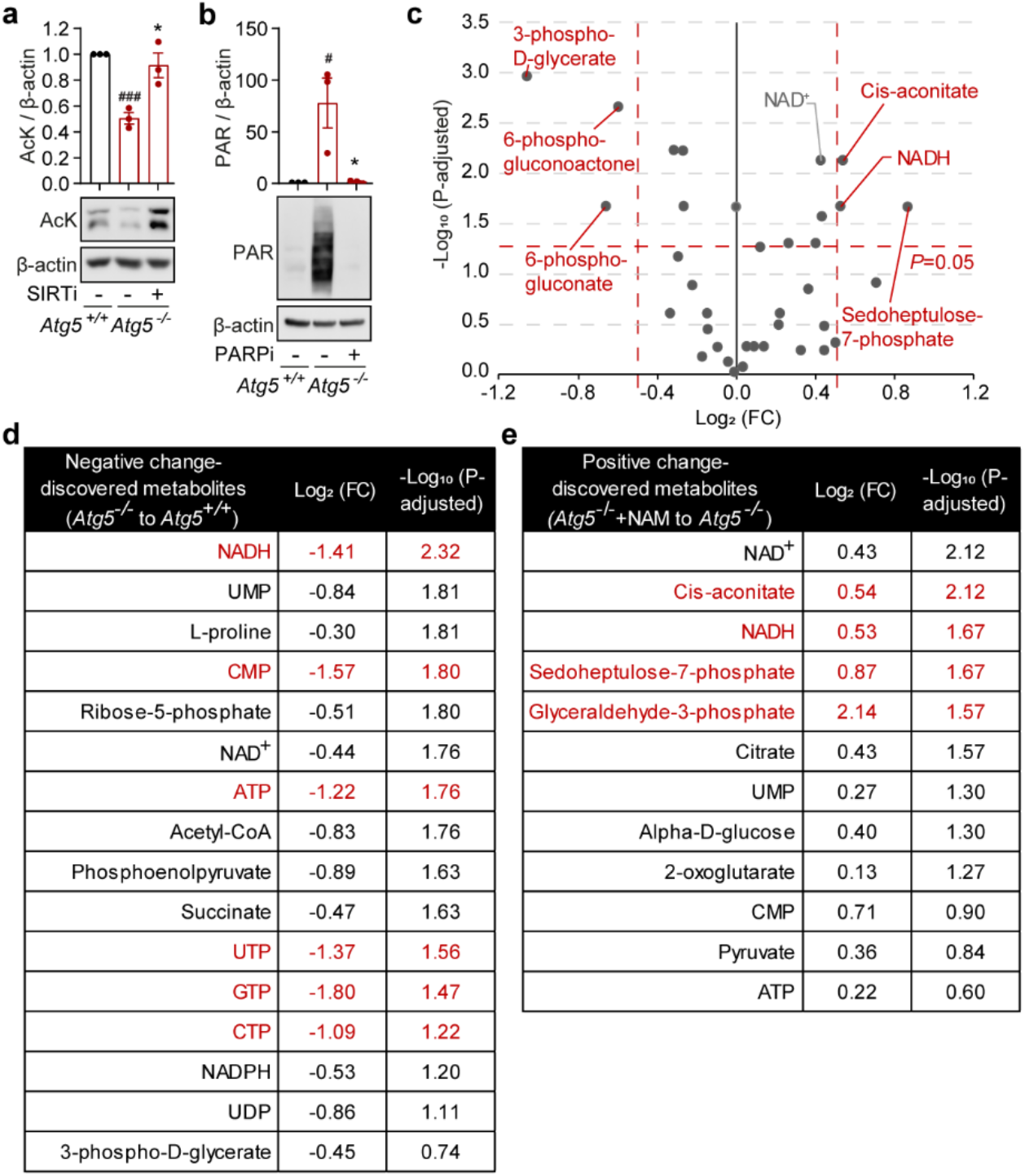
Levels of NAD(H) pools in *Atg5*^−/−^ MEFs correlate with NADase activities and cell viability. **a**, **b**, Immunoblot analyses with acetylated lysine (AcK) (**a**), poly(ADP-ribosylation) (PAR) (**b**) and β-actin antibodies (**a**, **b**) in *Atg5*^+/+^ and *Atg5*^−/−^ MEFs cultured in gal medium for 24 h treated with 20 μM sirtinol (SIRTi) (**a**), 10 μM olaparib (PARPi) (**b**) or solvent (DMSO) (**a**, **b**). **c**, Volcano plot representation of all analysed metabolites in a pairwise comparison of *Atg5*^−/−^+NAM to *Atg5*^−/−^ MEFs cultured in gal medium for 20 h. The significance cut-off was set to an adjusted *P*-value of 0.05 (−Log_10_(*P*-adjusted)>1,3) and a 1.4 fold-change (FC)(−0.51≥Log_2_(FC)≥0.49). Thresholds are shown as dashed red lines. **d**, **e**, Lists of ‘discovered’ metabolites that correlate with cell death/survival, i.e. depleted in a pairwise comparison of *Atg5*^−/−^ to *Atg5*^+/+^ MEFs and enriched in a pairwise comparison of *Atg5*^−/−^+NAM to *Atg5*^−/−^ MEFs (**e**). Significantly changing metabolites (−Log_10_(*P*-adjusted)>1.3, −1≥Log_2_(FC)≥1) (**d**) (−Log_10_(*P*-adjusted)>1.3, (−0.51≥Log_2_(FC)≥0.49) (**e**) are highlighted in red. Marked in bold are metabolites that change in correlation with *Atg5*^−/−^ MEF cellular survival. Data (**a**, **b**) are mean ± s.e.m. *P* values were calculated by unpaired two-tailed Student’s *t*-test (**a**,**b**) and the multiple *t*-test original FDR method of Benjamini and Hochberg (**c**, **d**, **e**) on three independent experiments. ^#^*P*<0.05, ^###^*P*< 0.001 (relative to *Atg5*^+/+^), \**P*<0.05 (relative to *Atg5*^−/−^).

**Extended Data Fig. 3:**
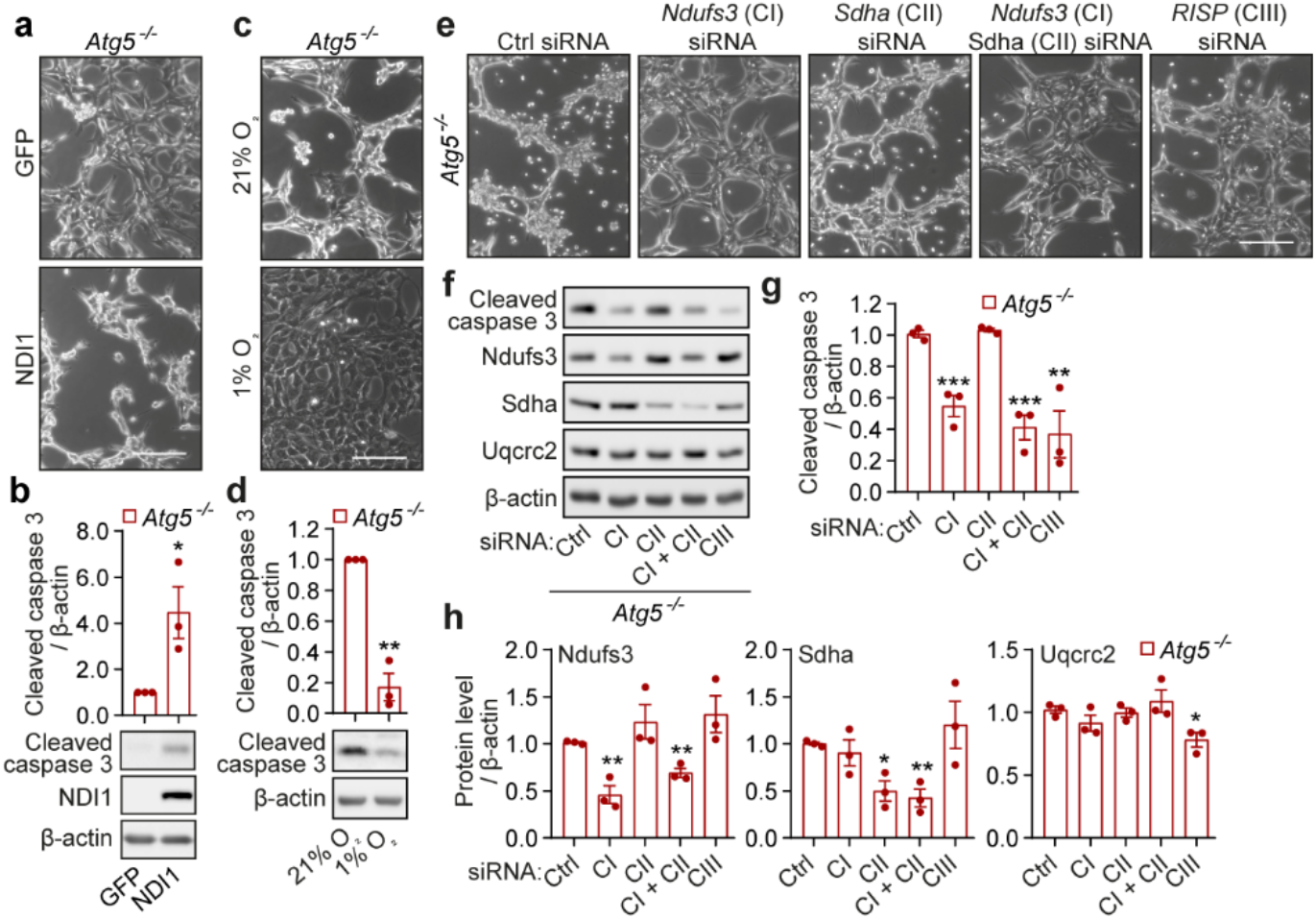
Mitochondrial respiration triggers apoptosis in *Atg5*^−/−^ MEFs. **a-d,** Phase-contrast images (**a**, **c**) and immunoblot analyses using cleaved caspase 3 (**b**, **d**), NDI1 (**b**) and β-actin antibodies (**b**, **d**) of *Atg5*^−/−^ MEFs transfected with NDI1-GFP (NDI1) or an empty plasmid (GFP) (**a**, **b**), or cultured in atmospheric oxygen (21% O_2_) or in hypoxia (1% O_2_) (**c, d**). **e**-**f**, Phase-contrast images (**e**) and immunoblot analyses with cleaved caspase 3, Ndufs3 (CI), Sdha (CII), Uqcrc2 (CIII) and β-actin antibodies (**f-h**) in *Atg5*^−/−^ MEFs transfected with control (ctrl), *Ndufs3*, *Sdha* and *RISP* siRNA individually and cultured in gal medium. Data (**b**, **d**, **g**, **h**) are mean ± s.e.m. *P* values were calculated by unpaired two-tailed Student’s *t*-test (**b**, **d**, **g**, **h**) on three independent experiments. \**P*<0.05, \*\**P*<0.01, \*\*\**P*<0.001 (relative to *Atg5*^−/−^). Scale bars, 200 μm (**a**, **c**, **e**).

**Extended Data Fig. 4:**
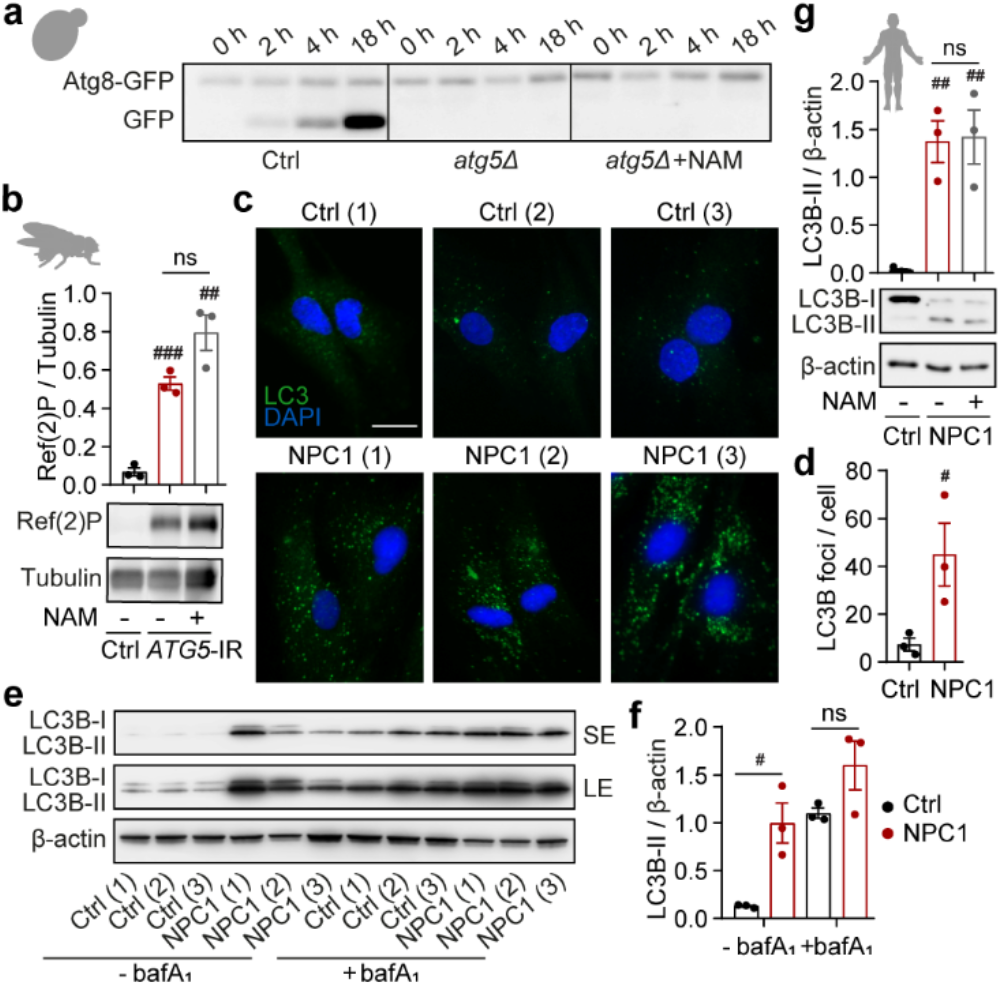
Investigation of autophagy impairment in yeast, fly and cell models. **a**, Immunoblot detection of Atg8-GFP cleavage as a readout of autophagy activation in ctrl and *atg5*Δ yeast strains at indicated time points. Displayed immunoblot is a representative of two independent experiments. **b**, Immunoblot analysis of whole-fly lysates from ctrl and *ATG5*-IR flies supplemented with 5 mM NAM or solvent (H_2_O) with Ref(2)P and tubulin antibodies. (*n*=10 flies per group). **c**, **d**, Immunofluorescence staining with LC3B antibody (**c**) and quantitation of intracellular LC3 puncta (**d**) in glucose-cultured healthy human volunteer (ctrl) and NPC1 patient-derived fibroblasts (NPC1) (*n*=3 cell lines per group). **e**, **f**, Immunoblot with LC3B and β-actin antibodies (**e**) and LC3-II quantitation in glucose-cultured ctrl and NPC1 fibroblasts with or without 400 nM bafilomycin A_1_ (bafA_1_) (**f**). Short exposure (SE) and long exposure (LE) of the same immunoblot are shown. **g**, Immunoblot analysis with LC3B and β-actin antibodies in ctrl and NPC1 patient fibroblasts sub-cultured in gal medium with or without 5 mM NAM. **c-g**, Details of control and NPC1 patient fibroblasts cell lines are listed in Supplementary Table 1. Data (**b**, **d**, **f**, **g**) are mean ± s.e.m. *P* values were calculated by unpaired two-tailed Student’s *t*-test (**b**, **d**, **f**, **g**) on two (**d**, **f**, **g**) or three (**b**) independent experiments. ^#^*P*<0.05, ^##^*P*< 0.01, ^###^*P*< 0.001 (relative to ctrl), ns – non-significant. Scale bar, 20 μm (**c**).

